# In situ Discovery of Immune Repertoire Reveals Antitumor Immunity and Therapeutic Antibodies

**DOI:** 10.64898/2026.08.11.744176

**Authors:** Haorui Zhang, Peiyu Wang, Yahui Zhao, Lukai Yang, Teng Xue, Linlin Liu, Yanping Zhao, Zongxu Zhang, Jiahao Ma, Baige Zeng, Peng Zhang, Cunyu Wang, Deng Pan, Zhidong Gao, Zhihua Liu, Zexian Zeng

## Abstract

Spatial transcriptomics offers a glimpse into the immunology of tissues. However, limitations in spatial transcriptomics preclude the detection of highly diverse, low-abundance, and previously unknown sequences, including immune repertoires and microbiota. Here, we introduce Archimap, a spatial transcriptomic platform that simultaneously profiles spatial transcriptomes, immune repertoires, and microbiota from formalin-fixed paraffin-embedded (FFPE) tissues. Using Archimap, we profile the spatial localization of TCRs, BCRs, and the microbiota landscape in archived clinical tissues at single-cell resolution. Through comprehensive benchmarking, we validate Archimap’s performance and fidelity. Archimap in situ assembles the immune complex and reconstructs the clonal evolution of antibodies. Together, Archimap shows the power of in situ discovery of functional immune repertoires for their antitumor immunity.

## Introduction

Adaptive immune repertoires and tissue-colonized microbiota are major determinants of human health and disease.^1-6^ B cells and T cells recognize antigens through highly diverse B cell receptor (BCR) and T cell receptor (TCR) repertoires generated by V(D)J recombination and somatic hypermutation (SHM), producing up to 10^13^∼10^14^ distinct receptor sequences.^7^ These receptor sequences provide a molecular record of immune-cell lineage and diversification.^8-13^ Likewise, microbial communities comprise evolutionarily diverse lineages that can be identified through 16S and 18S ribosomal RNA (rRNA) sequences.^6,14-16^ Together, immune repertoires and microbiota shape anti-tumor immunity, infection, and tissue homeostasis. Yet how these immune and microbial populations are spatially organized within tissues and how they interact with local microenvironments remain largely unknown.

Spatially organized immune repertoires provide unique opportunities to study tissue immunity and disease. Spatially organized and shared BCR and TCR clonotypes can help reveal antigen-specific immune responses, clonal expansion, and evolutionary trajectories within tissues. Beyond their value as lineage tracers, patient-derived immune repertoires have emerged as a source of therapeutic antibodies and TCR-T therapies.^17,18^ However, identifying functional receptors from millions of candidate clonotypes remains a major challenge and often relies on extensive screening with limited biological context.^19^ Spatial information could help prioritize clonotypes by linking them to disease-associated niches, cellular interactions, and functional immune states, thereby facilitating the discovery of therapeutic receptors.^10,12^ Similarly, spatial mapping of microbiota could reveal how microbial communities interact with local host immune systems and influence tumor progression and treatment response.^14^

Recent advances in spatial transcriptomics have transformed the study of tissue organization.^20^ A technology capable of resolving spatial immune repertoires and microbiota directly from archival FFPE samples would substantially expand access to patient cohorts and accelerate both mechanistic studies and therapeutic discovery. Here, we developed Archimap (Archival Immune Map), a platform that simultaneously captures spatial transcriptomes, paired immune repertoires, and microbiota from the same tissue sections. These results establish a framework for integrating spatial transcriptomes, immune repertoires, and microbiota within archival human tissues and facilitate the discovery of immune mechanisms and therapeutic candidates directly from clinical specimens.

## Results

### Archimap achieves high-fidelity immune repertoire, microbiota, and transcriptome profiling in FFPE tissues

The Archimap workflow is compatible with both freshly collected specimens and long-term FFPE archives. Following tissue sectioning, Archimap generates spatially resolved histological, transcriptomic, immune repertoire, and microbiome profiles from the same tissue section (**Figure 1A**). To benchmark Archimap, we profiled four treatment-naïve colorectal cancer (CRC) samples, named as POC(proof-of-concept)-01 to POC-04 (**Figure 1B and S1A**). Across these four samples, Archimap captured an average of 360,133 BCR counts and 7,979 TCR counts (**Figure S1B**), corresponding to 18,746 BCR clonotypes and 1,714 TCR clonotypes detected (**Figure S1C**), and to 70,966 B cells and 6,625 T cells detected per sample (**Figure S1D**). In parallel, Archimap captured an average of 6,461 microbiota-derived counts that could be mapped to microbial reference genomes (**Figure S1B**).

**Figure 1.**
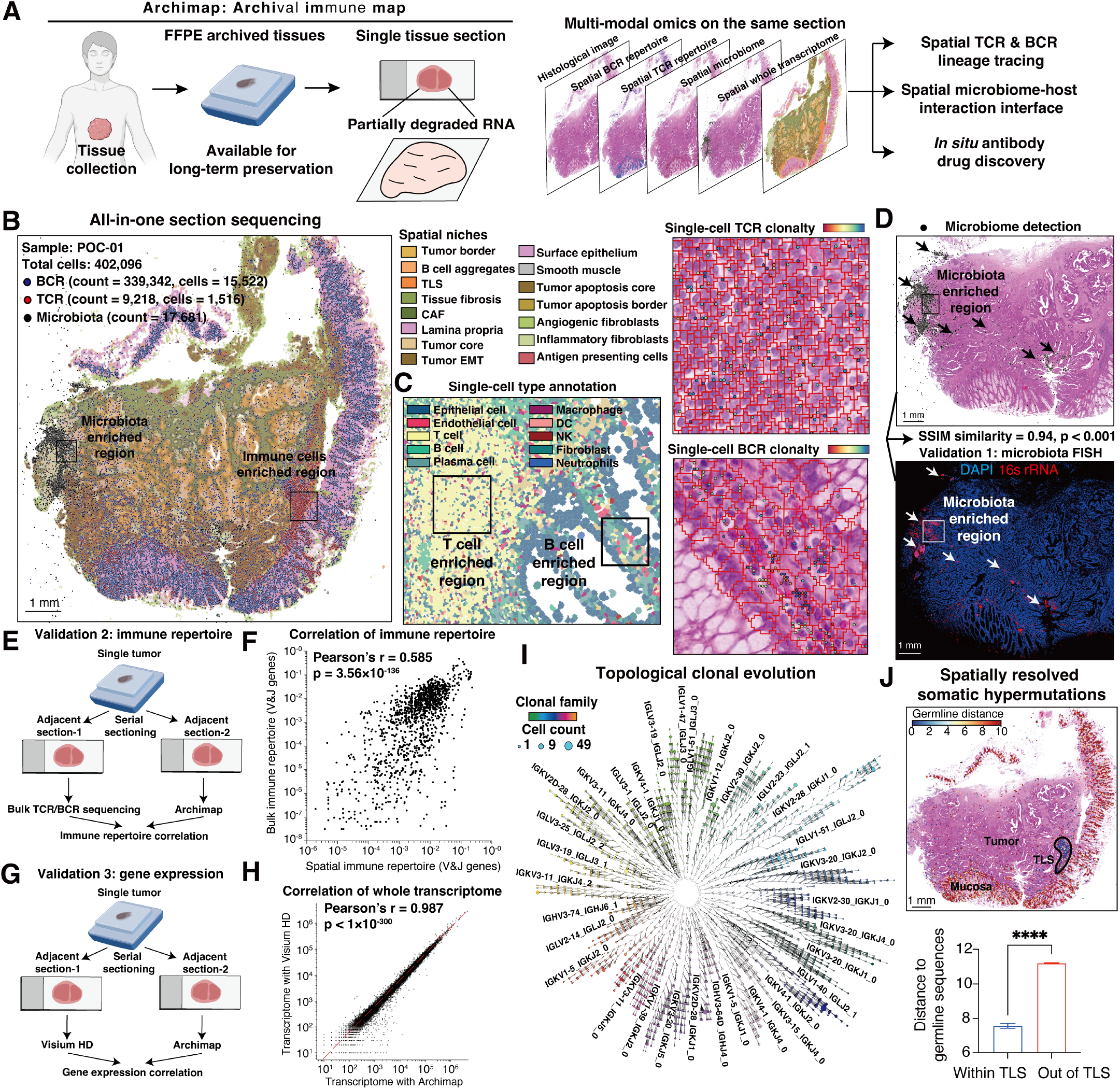
Archimap enables simultaneous spatial profiling of immune repertoires, microbiota, and transcriptomes in FFPE tissues. (**A**) Overview of the Archimap workflow for simultaneous spatial profiling of TCRs, BCRs, microbiota, and transcriptomes from FFPE tissues. (**B**) Representative Archimap dataset generated from an 11 x 11 mm FFPE tissue section, showing spatial niches, cell counts, immune clonotypes, and microbiota. (**C**) Single-cell annotation (left), and spatial distributions of single-cell TCR (right, upper) and BCR (right, lower) clonotypes in a magnified region of the tissue section shown in (B). (**D**) Validation of microbiota mapping by comparison with 16S rRNA FISH on an adjacent section. SSIM (structural similarity index measure) scores and corresponding P values are indicated. (**E**) Experimental design for validation of Archimap’s immune repertoire profiling. Adjacent tissue sections were profiled using traditional bulk TCR/BCR sequencing and Archimap. (**F**) Correlation of immune repertoire gene usage measurements comparing bulk TCR/BCR sequencing and Archimap. (**G**) Experimental design for validation of Archimap’s whole transcriptome profiling. Adjacent tissue sections were profiled using Visium HD and Archimap with a matched sequencing depth of 500 million reads. (**H**) Correlation of whole-transcriptome measurements comparing Visium HD and Archimap. (**I**) Topological visualization of the top expanded clonal families (n = 32) within the POC-01 tumor sample. (**J**) Spatial visualization (upper) and statistical analysis (lower) of the somatic hypermutation rate (measured by distance to germline sequences) for all clonal families, comparing regions within TLSs and outside TLSs.

We compared Archimap with the previously reported Spatial-VDJ platform.^10^ Notably, while Spatial-VDJ can only be performed for fresh-frozen tissues, Archimap was applied to archived FFPE samples, which generally contain more degraded and fragmented RNA. Despite this challenge, Archimap achieved 18.96-fold and 10.04-fold higher recoveries of BCR and TCR CDR3 counts per unit tissue area (**Figure S1E**).

To assess the spatial resolution of Archimap, we performed single-cell segmentation using Stardist,^21^ followed by cell-type annotation (**Figure 1C**). Distinct TCR and BCR clonotypes localized predominantly to regions enriched for T cells and B cells, respectively, and were concordant with both cell-type annotations and tissue morphology (**Figures 1C and S1F-S1G**). Consistently, Archimap-derived immune repertoire signals showed strong agreement with probe-hybridization-based measurements of BCR and TCR constant region (C region) expression, with Pearson correlations of 0.95 and 0.80 for BCRs and TCRs, respectively (**Figure S1H**).

To validate the resulting spatial distribution of the microbiota, we performed a serial sectioning experiment followed by 16S rRNA fluorescence in situ hybridization (FISH), a standard method for microbiota detection (**Figure 1D**).^15,16,22^ Archimap-derived microbiota signals exhibited strong concordance with FISH measurement (SSIM (structural similarity index measure) = 0.94, p < 0.001) (**Figure 1D**), demonstrating the high fidelity of spatial microbiota profiling in FFPE tissues.

We used an adjacent serial section to test the capture efficiency of immune repertoires. Adjacent sections from the four POC tumor samples were extracted for total RNA and were subjected to bulk TCR&BCR sequencing (**Figure 1E**). As a result, we found that both V and J gene usage frequency and the count of CDR3-determined clonotypes were significantly correlated with bulk sequencing as the standard (**Figures 1F and S1I-S1K**). Specifically, the V and J genes showed an overall Pearson correlation of 0.585 (p = 3.56×10^-136^) (**Figures 1F and S1I**), and the CDR3 clonotypes showed an overall Pearson correlation of 0.492 (p < 1×10^-300^) (**Figures S1J-S1K**).

**Figure S1.**
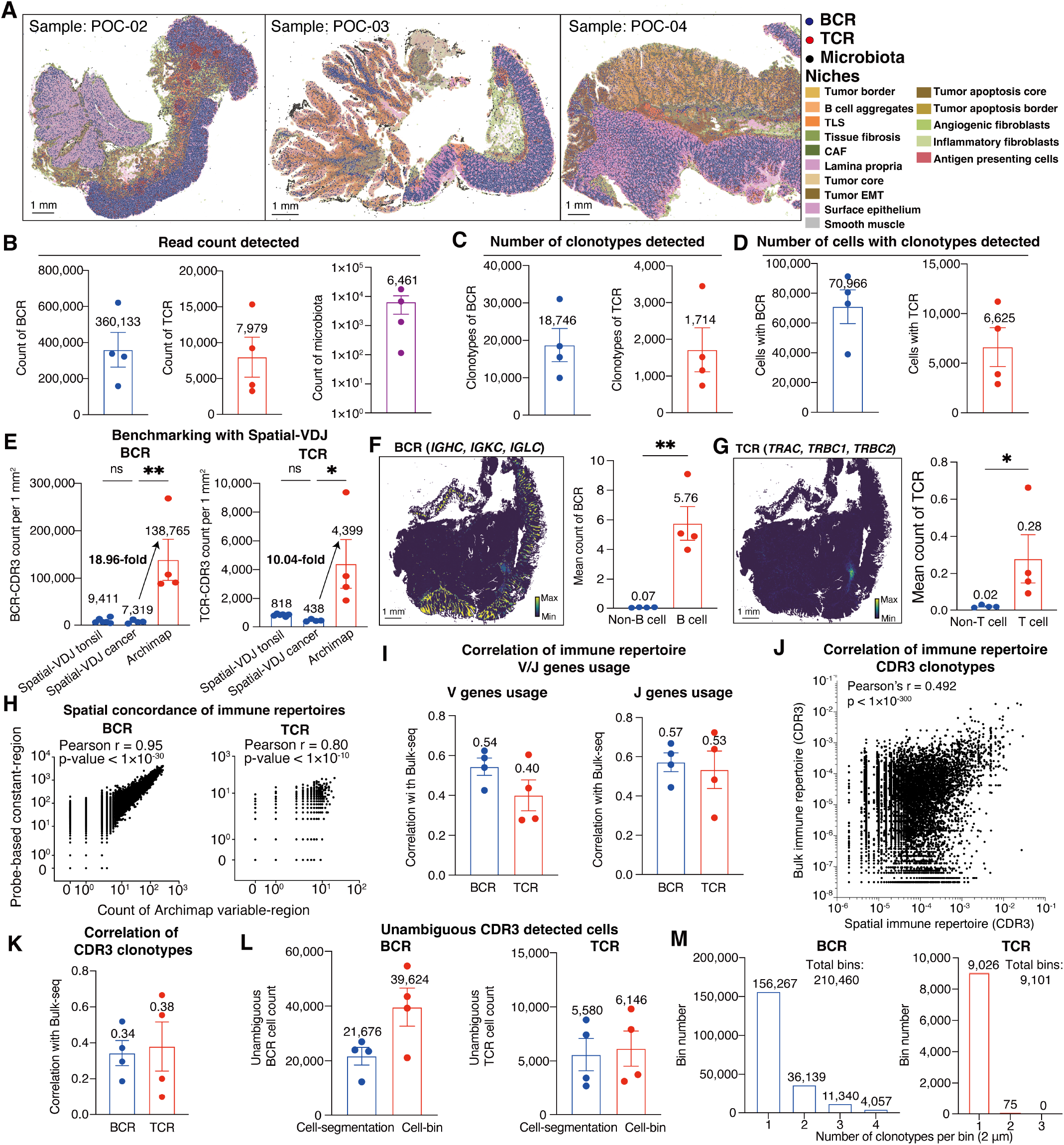
Benchmarking and validation of Archimap. (**A**) Spatial maps of three additional CRC tumor samples profiled by Archimap. (**B-D**) Quantification of Archimap detection across four samples, showing the read count of immune repertoire and microbiota (B), the number of detected clonotypes (C), and the number of TCR-BCR-positive cells (D). (**E**) Benchmarking of Archimap against Spatial-VDJ for BCR (left) and TCR (right), comparing the count of CDR3 detected, normalized to the tissue area of 1 mm^2^. (**F**) Spatial distribution of BCR repertoire abundance (left) and statistical comparison of BCR CDR3 counts between B-cell-enriched and non-B-cell regions (right). (**G**) Spatial distribution of TCR repertoire abundance (left) and statistical comparison of TCR CDR3 counts between T-cell-enriched and non-T-cell regions (right). (**H**) Spatial concordance between probe-based BCR and TCR constant-region expression and Archimap-derived variable-region abundance across 8 μm cell bins, supporting the spatial specificity of immune repertoire detection by Archimap. (**I**) Sample-wise correlations of V genes (left) and J genes (right) usage comparing bulk TCR/BCR sequencing and Archimap on adjacent sections. (**J-K**) Combined correlation (J) and sample-wise correlations (K) of immune repertoire at the CDR3 level comparing bulk TCR/BCR sequencing and Archimap on adjacent sections. (**L**) Number of segmented cells or cell-bins (8 μm) with unambiguous BCR (left) and TCR (right) detection in four tissue sections. (**M**) Assessment of spatial diffusion in Archimap-derived immune repertoires, with most BCR- and TCR-positive bins (2 μm) containing a single clonotype.

The third serial sectioning experiment was applied to evaluate whether Archimap introduces bias into whole-transcriptome profiling (**Figure 1G**). To this end, we performed serial sectioning of an additional CRC sample and generated matched datasets using either Archimap or the commercial Visium HD platform (**Figure 1G**). Whole-transcriptome expression profiles obtained by Archimap were highly concordant with those generated by Visium HD (Pearson r = 0.987, p < 10^-300^) (**Figure 1H**), indicating that targeted RT introduces minimal bias into transcriptome-wide measurements.

We next evaluated immune repertoire detection at both single-cell and cell-bin resolutions. Cell-bin analysis substantially increased the number of cells with unambiguous repertoire assignment, yielding averages of 39,624 BCR-positive and 6,164 TCR-positive cells at cell-bin level, and 21,676 B cells and 5,580 T cells at single-cell resolution, respectively (**Figure S1L**). Therefore, we used cell-bin resolution for subsequent analyses to maximize clonotype recovery while preserving spatial information. We further assessed the extent of spatial diffusion by quantifying clonotype redundancy per 2-μm spatial bin. Notably, 74.25% of BCR-positive bins and 99.18% of TCR-positive bins contained a single clonotype (**Figure S1M**), indicating minimal signal diffusion and high spatial specificity.

### Archimap in situ assembles BCR and TCR complexes and studies the topological clonal evolution

Immune receptors, BCRs and TCRs, are complexes formed with single chains. Specifically, BCRs are formed with heavy-chain and light-chain, while TCRs are formed with α-chains and β-chains. Assembly of these complexes at the tissue level remains challenging in two critical steps: the simultaneous capture of all CDRs,^23^ and the assembly of chains at high spatial resolution (**Figure S2A**).^10^ To this end, we asked whether Archimap could enable *in situ* reconstruction of immune receptors from FFPE tissues.

In the first step (**Figure S2A**), Archimap simultaneously captures the CDR1, CDR2, and CDR3 regions, enabling reconstruction of full-length antibody variable regions, which exhibit concordant spatial distributions across tissue regions (**Figure S2B**). Pairwise assembly rates between CDR regions exceeded 55% at 2 μm resolution (**Figure S2C**), and the average assembly rate for full-length Ig variable regions reached 28.43% at cell-bin resolution (**Figure S2D**).

In the second step (**Figure S2A**), Archimap simultaneously captures paired immune receptor chains, including antibody heavy-light chain pairs (IgH-IgL and IgH-IgK) and TCR αβ pairs (TRA–TRB), which also exhibited concordant spatial distributions (**Figure S2E**). At cell-bin resolution, pairing rates achieved 50% for BCR pairs and 23% for TCR pairs (**Figure S2F**).

We applied these *in situ*-assembled BCR complexes to study their topological evolution, represented by BCR somatic hypermutations in the CDR1, CDR2, and CDR3 regions upon antigen stimulation. Archimap captured single-base somatic hypermutations in these regions, thereby reconstructing the time-resolved lineage tree of the spatial immune repertoire (**Figures 1I and S2G**) and successfully mapping them to their spatial locations (**Figure S2H**). These reconstructed clonal trees showed spatially distinct distributions correlated with topological clonotype levels (**Figure S2I**), Ig class (**Figure S2J**), and spatial locations (**Figure S2J**), representing a lineage trajectory from priming, development, and maturation to terminal clonotypes (**Figures S2I-S2J**).

We validated our reconstruction of BCR topological evolution using the rule of antibody class switching, that is, the directed conversion of IgM to IgG and IgA (**Figure S2K**). As expected, over 75% of clonotypes successfully matched the class-switching rule, further validating the fidelity of the topological and spatial clonal trees (**Figure S2L**).

**Figure S2.**
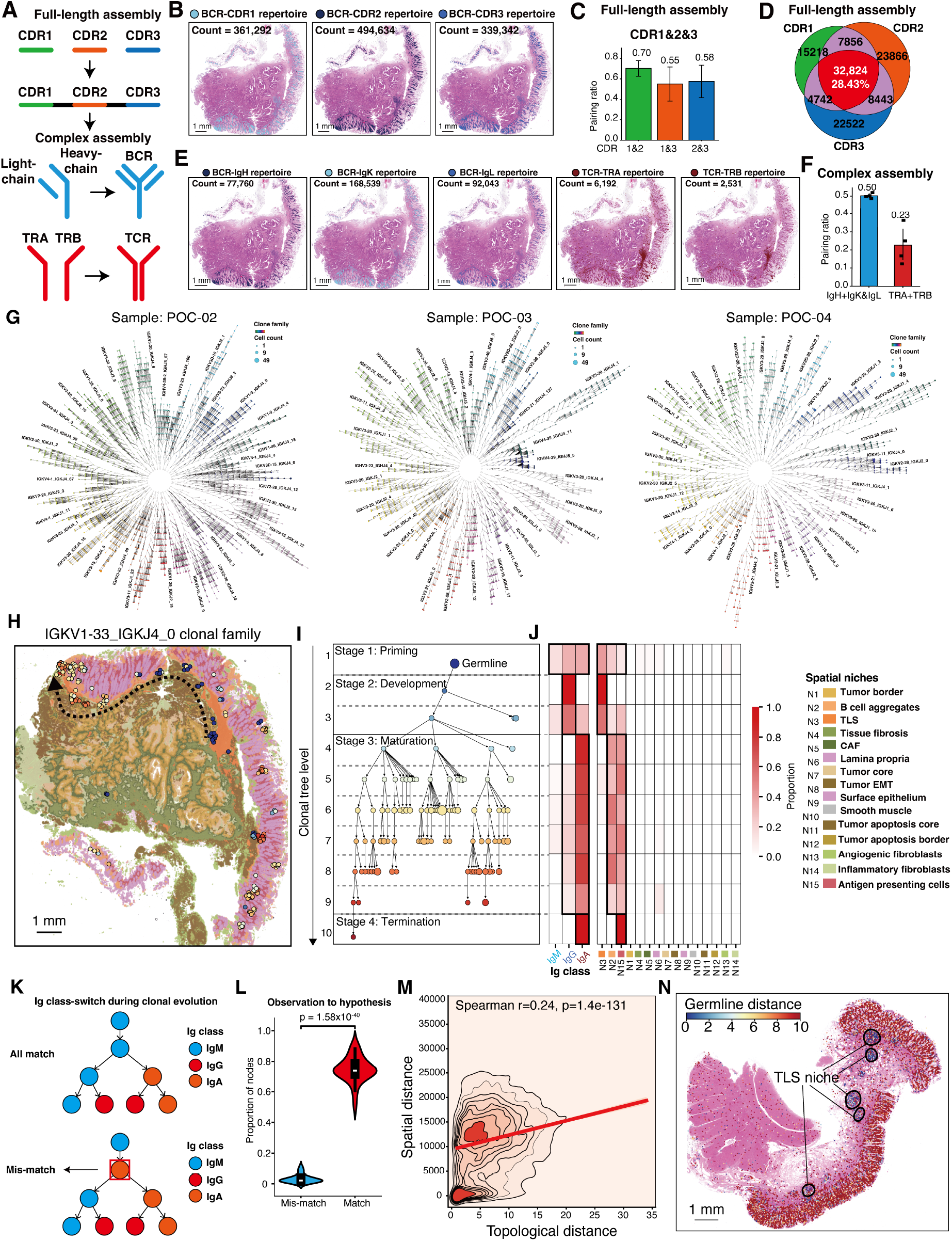
Immune receptor assembly and topological evolution. (**A**) The schematic of *in situ* immune complex assembly: the first step: pairing CDR1, CDR2, and CDR3 to build the full-length single chain, and the second step: pairing the light chain and heavy chain for BCR, and pairing the TRA and TRB for TCR. (**B**) Spatial distributions and density of BCR CDR1, CDR2, and CDR3 regions detected by Archimap. (**C**) Efficiency of *in situ* receptor assembly, including BCR CDR regions. (**D**) Assembly of full-length immune receptor chains at cell-bin (8 μm) resolution. Venn diagrams show the overlap of BCR CDR1, CDR2, and CDR3 regions. (**E**) Spatial distributions of IgH, IgK, and IgL repertoires across the tissue section. (**F**) Efficiency of *in situ* receptor assembly, including BCR heavy- and light-chains at cell-bin (8 μm) resolution.. (**G**) Topological visualization of the top expanded clonal families (n = 32) for POC-01, POC-02, and POC-03 tumor samples. (**H-J**) Spatial visualization of the representative clonal family (H), the topological evolution of clonotypes within the family (I), the Ig isotype expressed (J), and also the spatial localization of these different levels of clonotypes (J). (**K**) The schematic of Ig class-switching in the context of a match (upper) and a mismatch (lower), compared with previous reports. (**L**) Ratio of matched clonotype nodes compared with those mismatched clonotype nodes. (**M**) Correlation of clonal evolution depth with the spatial distances between each clonotype and the closest germline clonotype. (**N**) Spatial visualization of the germline distance of clonotypes detected within the TLS niche and other regions for POC-02 tumor samples.

Another piece of evidence supporting our results is the correlation between spatial distance and germline distance (**Figure S2M**). We found that as clonotypes evolved, the spatial proximity between each clonotype and its germline clonotype was significantly correlated (p = 1.4 × 10^^-131^) with the clonotype’s topological depth (**Figure S2M**). This evidence indicates that as BCRs undergo topological evolution, they also migrate to distant regions.

Next, we leveraged these topological evolutions as temporal references to investigate the potential origins of tissue-infiltrated clonotypes (**Figures 1N and S2N**). Indeed, we found significantly lower (p < 0.001) somatic hypermutations within the TLS-like niche across all clonal families (**Figures 1N and S2N**). These results suggest that TLS-like structures contribute to the stimulation of naïve clonotypes, which will then spatially infiltrate into tissue niches.

Together, these results establish Archimap as a scalable platform for integrated spatial profiling of immune repertoire, microbiota, and transcriptomes in archived clinical tissues, enabling in situ reconstruction of immune receptors at single-cell resolution.

## Discussion

Rapid advances in spatial transcriptomics have accelerated tissue-level biological discoveries.^24-26^ In this article, we developed Archimap, scalable for capturing clinical archived samples for their immune repertoires and microbiota. Archimap’s ability to discover immune repertoires in FFPE samples makes it scalable for the retrospective recovery of biological insights from samples archived for years, including previous clinical trials.

Archimap *in situ* captures the clonotypes and clonal families for TCRs and BCRs, and also somatic hypermutations for BCRs. Combining these features within a single section enables us to profile the spatial and topological evolution of BCRs. Topologically, clonal families differed in width and length, reflecting their bursts of somatic hypermutation. Further efforts should be made to identify whether BCR signaling or antigen stimulation determines these distinct topological patterns.

In the future, we anticipate that the *in situ* discovery of immune repertoires will contribute to both the deep dive into the tissue microenvironment for clinical patients, with such immune cell spatial tracing, and also the screening for potential therapeutics.

